# Dominance asymmetries shape vocal exchanges in meerkats

**DOI:** 10.1101/2024.11.25.624808

**Authors:** Vlad Demartsev, Gabriella Gall, Ariana Strandburg-Peshkin, Marta B Manser

## Abstract

Physical grooming is often used to maintain social bonds in animal groups, but opportunities for grooming each group member may be limited. Vocal exchanges have been proposed as an alternative way to sustain social ties without requiring physical proximity. To explore the link between vocal exchanges and social relationships, we examined the effect of dominance and dyadic bond strength of the interacting partners on the intensity of vocal interaction in meerkats (Suricata suricatta). Focusing on ‘sunning calls’, socially facilitated vocalisations produced when meerkats warm themselves in the sun, we conducted experiments where we played sunning calls to individuals and recorded their responses. Overall, subordinate individuals showed an increased call rate when responding to calls from dominants. Tie strength showed a weak trend that differed across dyad types. These results support the notion of meerkat sunning calls having a social regulatory function, possibly as a bonding or appeasing signal, mainly directed to dominant individuals. Our results reflect the dichotomy between the subordinate and dominant individuals and underscore the strategic nature of social relationships within a despotic society. Engaging in continuous reciprocal interactions can signify commitment, promoting tolerance and cooperation. Investing in such interaction with influential dominants could lower the chances of aggression and would fit the ‘vocal grooming’ functionality of such call exchanges. Further work establishing causal links between vocal interaction patterns and long-term relationship outcomes will be crucial for understanding the scope of ‘vocal grooming’ as a complement or alternative to physical affiliation across species.

**LAY SUMMARY:** Meerkats use more than grooming to manage their social relationships—they also exchange short calls while warming in the sun. Playback of these “sunning calls” to wild meerkats led to increased calling by listeners, particularly when subordinates heard dominants. Females responded somewhat more than males, and social bond strength had a modest influence on calling patterns. Such vocal exchanges may function as “vocal grooming,” helping maintain relationships at a distance.

## INTRODUCTION

How group-living animals maintain stable social relationships is a central question in behavioural ecology. Social bonds can be reinforced through physical interactions such as grooming, but opportunities for grooming may be limited in large groups or when individuals are spatially dispersed. Robin Dunbar’s (1998) social bonding hypothesis suggests that vocal interactions among group members can complement and perhaps partially substitute the social function of physical grooming. Frequently, the tendency to respond to vocal signals of conspecifics can be correlated with social affinity. In many social species, strongly bonded individuals engage in both mutual physical grooming and vocal interactions more frequently than those with weaker social bonds (Arlet et al., 2015; Kulahci et al., 2015). For example, dyads of male chimpanzees (*Pan troglodytes schweinfurthii*) were more likely to engage in other affiliative behaviours such as grooming on days when they participated in pant-hoot displays, suggesting that joint vocal activity might represent bonding (Fedurek et al., 2013). Yet the relationship is not always positive: in bottlenose dolphins (*Tursiops aduncus)*, weakly bonded individuals exchange more signature whistles than strongly bonded pairs, who rely more on physical affiliation (Chereskin et al., 2022). Such variation suggests different strategies for maintaining existing social ties or improving the strength and stability of weak ties, both of which have fitness implications for group-living animals (Lusseau et al., 2006). From the perspective of the “vocal grooming” hypothesis: (i) if vocal exchanges substitute for grooming, they should be more frequent where grooming opportunities are constrained (e.g. with weakly bonded partners); (ii) if they reinforce strong bonds, they should be more frequent between closely affiliated individuals. In either case, the distribution of vocal exchanges should reflect the social relationships among group members.

Most of the existing evidence linking the social function of grooming and vocal exchange behaviours is derived from studies of primates (Arlet et al., 2015; Kulahci et al., 2015; McComb and Semple, 2005). However, recently, the association of primary bonds with affiliative physical interactions and of secondary bonds with vocal exchanges have been reported in bottlenose dolphins (Chereskin et al., 2022). This raises the question of how vocal interaction intensities in other social mammals map onto social asymmetries. Here, we investigated this question in meerkats (*Suricatta suricata)*, a cooperatively breeding mongoose species that lives in highly cohesive groups (Clutton-Brock and Manser, 2016). As the first step towards determining social regulatory function consistent with the vocal grooming hypothesis, we examined how both dominance status and dyadic tie strength influence individuals’ vocal responses to calls from different partners.

Meerkats show a despotic dominance hierarchy with a dominant breeding pair and subordinate helpers (Clutton-Brock and Manser, 2016). Within the subordinate class, the hierarchy is less obvious: age and body weight largely explain the dominance relationships among subordinate females (Thavarajah et al., 2014), however, overall dominance-related associations among subordinates are infrequent (Madden et al., 2011). Intra-group social interaction and cooperative behaviour patterns are also not associated with kinship (Clutton-Brock et al., 2001; Madden et al., 2012) and groups show no sexual segregation during diurnal activities (Gall and Manser, 2018). While conflict among meerkats is most extensive within the same sex, such intra-sex conflicts are low during the non-breeding season (Kutsukake and Clutton-Brock, 2006a). Instead, meerkats cooperate and coordinate their actions with all group members on a day-to-day basis (Gall and Manser, 2017). However, the dominant-subordinate asymmetry has been repeatedly shown to influence social relationships (Kutsukake and Clutton-Brock, 2006a; Kutsukake and Clutton-Brock, 2008a) in both reproductive and non-reproductive contexts (Kutsukake and Clutton-Brock, 2008a; Madden et al., 2009).

Physical grooming is one of multiple aspects of meerkat behaviour where a dominance dichotomy is evident (Madden et al., 2009). Adult subordinates were more likely to groom the dominant female than vice versa (Kutsukake and Clutton-Brock, 2006b) and dominant males received more grooming from other group members while providing less (Kutsukake and Clutton-Brock, 2010). The only fully reciprocal grooming relationship among adult individuals exists between the dominant male and female. As group size increases, grooming networks become sparser, suggesting that individuals are limited in the number of grooming partners (Madden et al., 2009). The duration of grooming by a dominant male decreases in larger groups (Kutsukake and Clutton-Brock, 2010), in agreement with Dunbar’s hypothesis (Dunbar, 1998). However, subordinates spend more time grooming the dominant male in larger groups, suggesting a higher investment in bond maintenance, perhaps due to stronger competition (Kutsukake and Clutton-Brock, 2010).

Thus, allogrooming in meerkats is linked to the regulation of social relationships via appeasement of the dominant individuals (Kutsukake and Clutton-Brock, 2006b; Madden and Clutton-Brock, 2009). Given the social organisation of meerkat groups (Madden et al., 2009) and their reliance on vocalisations for social monitoring (Demartsev et al., 2024; Reber et al., 2013) we hypothesised that meerkats’ vocal exchange patterns would reflect their social relationships and could serve as prosocial behaviour (behavior that produce benefits to others, Burkart et al., 2009), similarly to allogrooming. While grooming provides direct hygienic benefits, vocal exchanges may provide social benefits by reducing tension, lowering the likelihood of aggression, and promoting coordination.

A candidate vocalisation that we explored in this regard is the “sunning call”; a relatively soft (50–55 dB at ∼0.5 m, *unpublished data*), short and tonal call produced mainly in the morning after a group has emerged from its sleeping burrow and spends up to an hour warming up in the sun (Demartsev et al., 2018). Sunning calls are socially facilitated and only produced when other individuals are present or have been heard calling. Additionally, individuals avoid overlapping the calls of their group mates and call in a turn-taking pattern (Demartsev et al., 2018) suggesting a call-and-response interaction rather than a group chorusing behaviour. During the morning sunning period, there are relatively few physical social interactions between the individuals. However, on cloudy or windy mornings, when meerkats are unable to sun effectively, they often engage in grooming and play behaviour (VD, GG, ASP, MM personal observations), potentially indicating the time after emergence from the sleeping burrows as a “social time” aimed at reaffirming bonds before initiation of group travel and foraging. Acoustically, sunning calls resemble sentinel calls (Rauber and Manser, 2017), short note calls produced while running (Demartsev et al., 2024; Manser, 1998) and submission calls (Manser, 1998). However, while running short notes and submission calls are produced in long and rapid sequences, sunning calls are emitted either as a single note or in short (mostly double or triple note) bursts (Demartsev et al., 2018).

Sunning calls are produced during low-conflict, stationary sunning behaviour, and are structurally similar to submission calls. We therefore predicted that sunning call exchanges may be a form of social regulation influenced by the relationships between callers and receivers. Drawing parallels with grooming and as physical grooming in meerkats shows a pronounced dominance-related asymmetry, we predicted that subordinates would show increased responsiveness to the calls of dominant individuals. To test this prediction, we quantified the context-dependent social association between group members during the sunning period using a network analysis based on spatial proximity. In parallel, we conducted a series of playback experiments in which individuals were presented with sunning calls from different group members, allowing us to assess how both dominance status and dyadic bond strength influenced vocal responses.

## METHODS

We collected the data for this study at the Kalahari Research Centre (KRC), Northern Cape, South Africa (Clutton-Brock et al., 1998a) in June-August of 2019 and 2021. The individually marked meerkat population of the Kalahari Meerkat Project (KMP) on site is fully habituated to human presence, including behavioural observations and audio recordings from < 1m. All field procedures were performed according to existing protocols and were subject to the ethical regulations of the University of Pretoria, South Africa (permit: EC047-16) and the Northern Cape Department of Environment and Nature Conservation (permit: FAUNA 1020/2016).

### Social association

As a part of long-term data collection at the KMP, meerkat groups were continuously monitored with individual and behavioural data collected by a team of trained volunteer researchers. Through this dataset, we had access to information on age, sex and dominance status of all animals included in the study. To estimate social association within meerkat groups, we collected data on the physical proximity of individuals during sunning behaviour. Each morning, we observed the selected group of meerkats during their emergence from the sleeping burrow. In 2019, we observed four groups on average 7.25 (range 5-9) mornings, and in 2021, we observed three groups for 9.3 (range 8-12) mornings (see Table S1 for detailed group composition). To avoid unbalanced sampling due to the variable duration of the daily sunning period (Demartsev et al., 2018), we performed one full group scan per day. As soon as all group members emerged from the sleeping burrow and started sunning (sitting or standing on hind legs facing the sun with the ventral side of the body), we documented the identity and association of all individuals. Specifically, we considered all individuals observed within 1 m of each other as being together. At the start of the sunning session, the animals are usually static, and there is little rearrangement. We did not record subsequent movements and changes of positions that mostly occur shortly before groups depart from the burrow area and start foraging.

From the proximity data collected while sunning, we generated weighted, undirected networks. In these, each group member was represented as a node, and the edges between individuals were calculated as the number of times two individuals were observed sunning together, divided by the number of total observation days (all individuals were present in all days). It has been previously discussed that constructing social networks using small sample sizes can lead to misrepresentation of the strengths of individual social relationships (Lusseau et al., 2008). To estimate social tie uncertainty, we performed a parametric bootstrap (Ferrando et al., 2022; Tibshirani and Efron, 1993) for all constructed networks: For each dyad, we first calculated the observed proportion of days they were seen together (number of “together” days divided by total observation days). This proportion, P_obs_, was treated as the underlying daily probability of association for that dyad. We then modelled the daily encounter history as a sequence of n independent Bernoulli trials (where n is the number of days observed), each with probability P_obs_ of being “together”. More specifically, for each bootstrap replicate, we simulated a new daily history for the dyad by drawing n binary outcomes from a binomial distribution with parameters n=1 and p=P_obs_ for each day. From this simulated history, we recalculated the dyad’s tie strength. Repeating this process across 1000 replicates produced a sampling distribution of tie strengths for each dyad, from which we extracted the 2.5th and 97.5th percentiles as the 95% confidence interval. These per-dyad tie-strength distributions were later incorporated into the statistical analysis.

### Playback experiments

Before the playback trials, we recorded individual sunning vocalisations of all group members older than 6 months. For the audio recordings, we used a Marantz PMD-661 digital recorder (Marantz, Japan) with a sampling rate of 44.1 KHz, 16-bit and a directional Sennheiser ME66 microphone with K6 power module (Sennheiser electronic GmbH, Germany) placed on a telescopic boom pole and held at ∼30 cm from the focal animal. We examined the recorded files in Avisoft-SASLab Pro (Avisoft Bioacoustics e.K, Germany) and selected 30 to 60-second segments with clearly recognisable sunning calls, low background noise and no additional meerkat vocalisations. We constructed the playback tracks by adding a leading 30-second background noise segment as a “control phase” and an additional trailing 30-second segment of background noise as a “recovery phase”, resulting in 90 to 120-second tracks (ESM, Audio S1, Fig. S3). We used each track in up to three playback trials, with individual animals exposed to any specific track only once (ESM, Table 1).

**Table 1:** Output for loglinear GLMM model. Standardized tie length was the response variable. Dominance/Age dyad was the explanatory variable, with the Dominant-Dominant dyad as the reference category. The subordinate age classes are: **Adult** > 2 years., 2 > **Yearling** > 1 year, 1 > **Sub-Adult** > 0.5 years, the dominant male or female individuals are: **Dominant**. Group was included as a random effect. Significant effects are marked in bold. σ^2^ = 0.20, N Groups = 6, Observations = 525, Marginal R^2^ = 0.093.

|  | *Predictors* | *Estimates* | *CI* | *p* |
| --- | --- | --- | --- | --- |
| **Dominance/Age dyad** | (Intercept) | 0.36 | 0.24 – 0.53 | **<0.001** |
|  | Adult_Dominant | 0.60 | 0.41 – 0.90 | **0.012** |
|  | Adult_Adult | 0.57 | 0.37 – 0.87 | **0.010** |
|  | Adult_Yearling | 0.68 | 0.47 – 0.98 | **0.037** |
|  | Adult_Sub-Adult | 0.68 | 0.46 – 0.99 | **0.045** |
|  | Yearling_Dominant | 0.81 | 0.56 – 1.17 | 0.258 |
|  | Yearling_Yearling | 0.86 | 0.60 – 1.23 | 0.405 |
|  | Yearling_Sub-Adult | 0.86 | 0.60 – 1.24 | 0.425 |
|  | Sub-Adult_Dominant | 0.80 | 0.55 – 1.18 | 0.262 |
|  | Sub-Adult_Sub-Adult | 1.14 | 0.77 – 1.69 | 0.510 |
| **Zero inflated model** | (Intercept) | 0.24 | 0.20 – 0.30 | <0.001 |

We played the tracks using a BRV-1 Bluetooth speaker (Braven, Australia), frequency response 180Hz to 13kHz ± 3dB, positioned on the ground, in an inconspicuous spot behind a shrub or a sand mound and at a 0.5-1 m distance from a focal sunning meerkat. Using an Android mobile device, Nokia 7.1 (Nokia Corporation, Finland) paired to the speaker via Bluetooth, we activated the playback. Meerkats targeted by the playback were all > 6 months old and sunning at > 1.5 m distance from their nearest neighbours. On a given morning, no more than three trials were performed, and no individual was targeted as a focal receiver more than once. We kept a minimal interval of 15 minutes between consecutive trials and a minimal interval of 48 hours between trial days performed in the same group. Throughout the playback, we recorded the vocalisations of the focal individual and noted all behavioural responses. If during the payback the focal individual moved away or additional meerkats entered the 1.5 m radius, we terminated the trial and omitted it from the analysis.

For each trial, we documented the identities of both the individual whose sunning calls we used in the playback track (sender) and the focal individual (receiver). Given the pervasive role of dominance in meerkat social relationships, we constructed a “dominance dyad” variable, reflecting the dominance statuses of the sender and the receiver in each trial (e.g., in a Dominant→Subordinate trial: the calls of a Dominant individual (Clutton-Brock et al., 1998b) in the group were played to a Subordinate receiver). In total we performed 180 trials with 21 trials excluded from subsequent analyses due to: noisy recording conditions (5); conspecifics approaching the focal individual or the playback speaker (2); focal individual moving away during the experiment (8); alarm calls (2), errors in assigning experiment time markers (4). The 2019 analysed dataset consisted of 81 playbacks, targeting 28 meerkats in four groups with an average of 20.3 trials per group (range 12-30). The 2021 analysed dataset consisted of 78 trials in three groups, targeting 23 meerkats with an average of 26 trials per group (range 15-35). Our playback dataset sample sizes were: Dominant→Dominant x2, Dominant→Subordinate x28, Subordinate→Dominant x20, Subordinate→Subordinate x111. As the Dominant→Dominant dyad type was strongly underrepresented, we excluded these trials from subsequent analysis.

### Audio processing and statistical analysis

We inspected the recorded files in Avisoft-SASLab Pro and manually labelled both the recorded playback calls and the calls emitted by the receiver in the three phases of the trial (control, stimulus, recovery) using Avisoft cursors. For each of the playback phases, we calculated the call rate by dividing the number of calls emitted by the receiver by the duration of the playback phase. To control for contextual and individual variation in basal call rates, for each trial we subtracted the call rate in the control phase from the call rate in the stimulus and recovery phases. The resulting Δ call rate values represent an individual change in calling behaviour between the phases of a given playback trial, thus accounting for unobserved effects that could affect individuals’ basal call rate (e.g. calls of remote neighbours).

To test for differences in age and dominance-dependent association while sunning we fit a zero-inflated generalized linear mixed model (GLMM) with gamma distribution (ziGamma) with the tie-weights as the response variable and paired age-dominance relationship as an explanatory variable. Group identity was fitted as a random effect.

To test whether the sunning call playbacks affected individual call rates and thus whether call rate changed in different phases of the playback experiments, we fit pairwise Wilcoxon tests and controlled for multiple testing using a Benjamini and Hochberg (1995) correction. This was done, as the data did violate multiple assumptions of GLMMs and we thus decided to use a non-parametric test.

To test for the effects of sender - receiver dominance (dominance dyad), relationship (tie-strength), receiver sex and their interactions, we designed a multiple imputation (MI) procedure (Noble and Nakagawa, 2021). We used the full uncertainty distribution of the dyadic tie strengths from parametric bootstrap draws to generate 1000 independent datasets. We then fitted the same mixed-effects model to each imputed dataset. In the model, we specified Δcall_rate as the response, tie strength, dominance dyad and receiver sex as main effects. We included two interaction terms of tie strength with dominance dyad and with receiver sex. To account for repeated measures, we included random intercepts for caller identity and receiver identity nested within the group_year.

We summarised model effects by pooling coefficients across imputations (Schomaker and Heumann, 2018) and reporting empirical 95% quantile intervals of the coefficient draws, thereby capturing both model and imputation uncertainty. To assess the strength of evidence for each parameter, we calculated posterior sign probabilities (*P*(β > 0), *P*(β < 0)) as the proportion of coefficient draws greater or less than zero, respectively. We derived post-hoc slope contrasts between dyad types directly from the pooled draws across all 1,000 imputations, ensuring that both model and imputation uncertainty were incorporated into inference. We performed model diagnostics on representative imputation by examining residual distributions, residual–fitted value plots, and random effect estimates to confirm that model assumptions were met. Coefficient estimates were consistent across imputations, indicating stability of the MI procedure. DHARMa residual diagnostics on a representative imputation indicated approximately uniform residuals with a slight deviation, no evidence of over/under-dispersion, and no zero-inflation.

All statistical analyses were conducted in R 4.0.3 and 4.3.1 (R Development Core Team, 2020). We used mixed-effects models (Bates et al., 2015; Brooks et al., 2017) to account for repeated testing of the same receivers. Tests were two-tailed and considered significant at *P <* 0.05. For the linear model analyses, we checked that the model assumptions (normality of error, homogeneity of variance and absence of multicollinearity) were satisfied, using DHARMa (Hartig, 2017). We used the “phia” (De Rosario-Martinez et al., 2015) and “emmeans” packages (Lenth et al., 2021) for post-hoc analysis and calculation of p-values from the mixed effects models.

## RESULTS

We found that during sunning, the subordinate adult individuals (> 2 years) in the group had relatively weak associations with the two other subordinate age classes (yearling and sub-adults). Subordinate adults also had weak associations with dominant individuals of the group. The younger age classes (sub-adults and yearlings), on the other hand, were frequently sunning together with one another, and together with the dominant individuals, resulting in strong association (Fig.1, Table 1).

**Figure 1:**
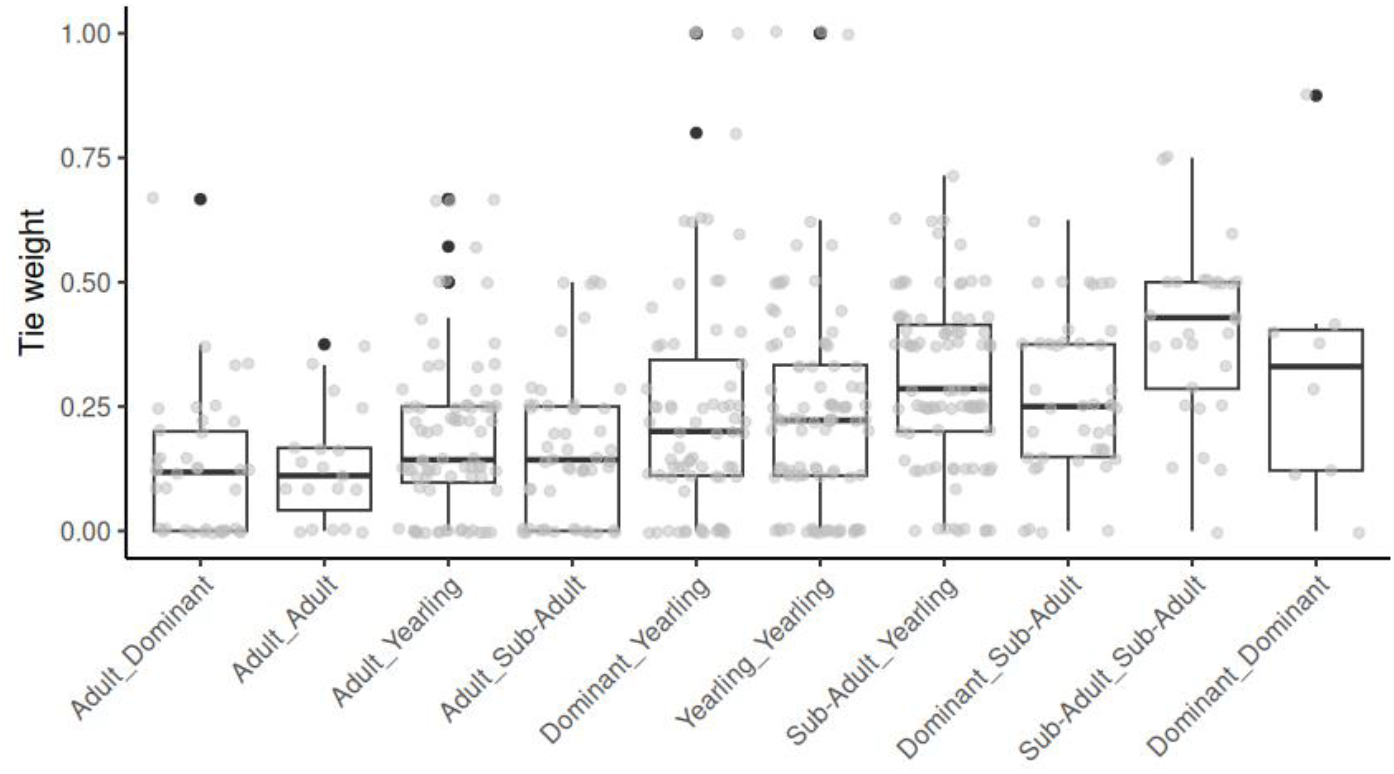
The distribution of tie strengths (y-axis) within Dominance/Age dyads (x-axis) in meerkat groups during sunning. Each category denotes the tie strength between pairs of individuals of a Dominance/ Age combination according to subordinate age classes: **Adult** > 2 years., 2 > **Yearling** > 1 year, 1 > **Sub-Adult** > 0.5 years, and the dominant male or female individuals: **Dominant**.

Examining the overall effect of sunning call playbacks on meerkat vocal behaviour, we found a significantly increased call rate of receivers in the trial phase (Wilcoxon Test: χ^2^ = 64.79, df = 2, P < 0.001, Figure 2, Table S2). Specifically, receivers increased their call rate during the stimulus phase compared to the control (P < 0.001) and the recovery phase (P = 0.046). Similarly, the call rate remained higher in the recovery phase compared to the control phase (P = 0.036).

**Figure 2:**
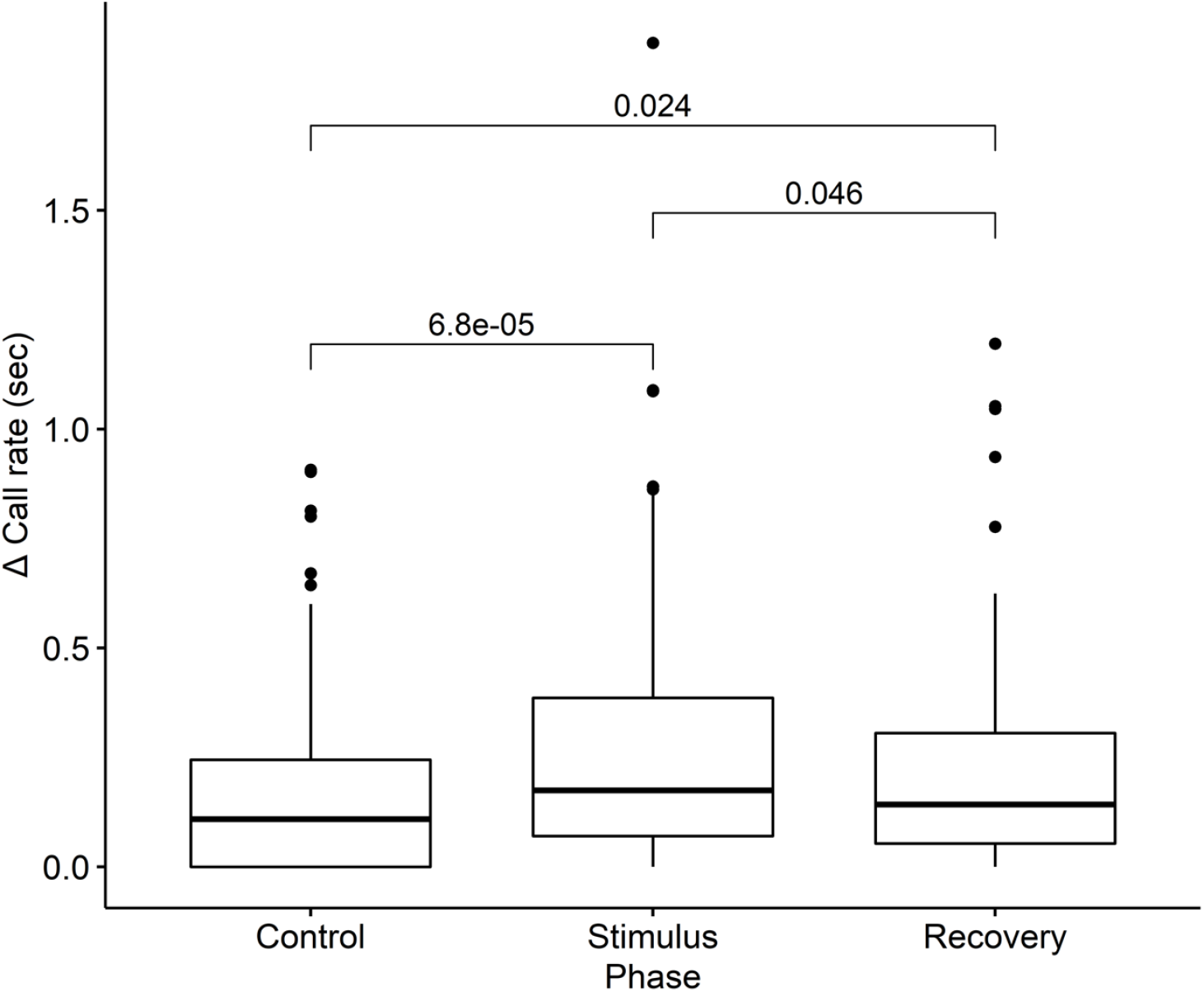
Receiver call rate in the three phases of the playbacks. In comparison with the Control, the call rate is significantly increased during the stimulus phase. The call rate during the Recovery phase is significantly lower in comparison to the Stimulus but significantly higher in comparison to the Control phase (Wilcoxon paired test).

Testing the magnitude of receiver’s vocal responses in detail, we found clear evidence for differences in call rate across dominance pairings and receiver sex (Table 2). Subordinate → Dominant and Subordinate → Subordinate dyads showed reduced responder call rates relative to Dominant → Subordinate pairs. Receiver females exhibited a small but consistent increase in call rate compared to males. For the effects of tie strength, overall slopes were estimated with wide uncertainty, and none differed clearly from zero. However, interaction terms revealed that the effect of tie strength varied across dominance pairings. In Dominant → Subordinate dyads, the tie-strength slope was negative in 92% of the draws and slightly positive in 76% of Subordinate → Subordinate dyads draws (Fig. 3, Table 2). Consistent with this pattern, the Dominant→Subordinate slope exceeded Subordinate→Subordinate slope in 94% of draws, although this contrast was not significant in the two-sided bootstrap test (p = 0.11; Table 2).

**Table 2:**
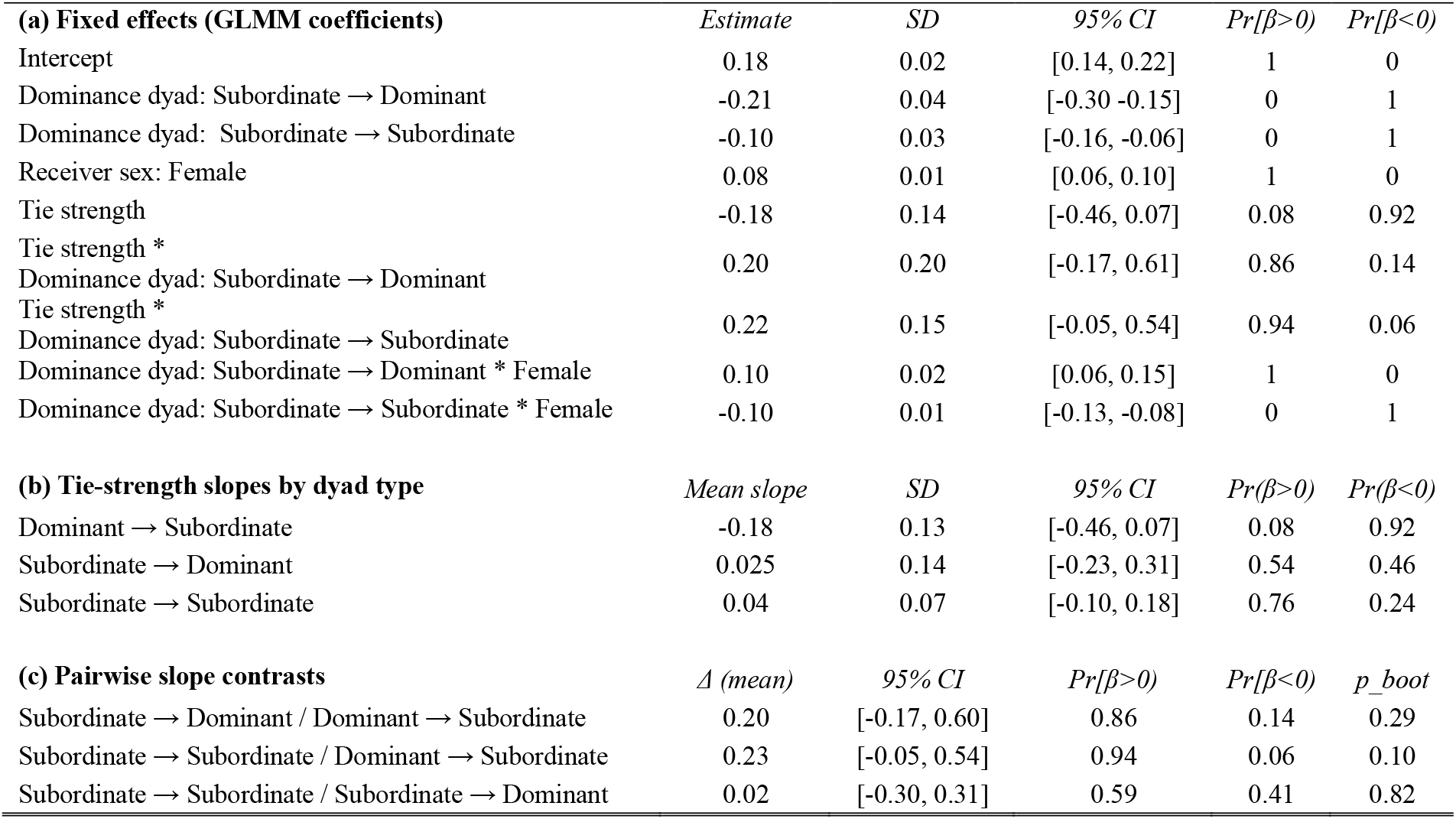
Results of mixed-effects models testing the effect of dominance pairing, tie strength, and receiver sex on meerkat vocal responses. (**a**) Regression coefficients from the full model. Estimates are MI-pooled means, with 95% quantile-based confidence intervals (CI) and the probability that each coefficient is greater than zero (Pr(β>0)) or less than zero (Pr(β<0)). The intercept corresponds to Dominant → Subordinate pairings with male receivers. (**b**) Predicted slope of the relationship between tie strength and receiver call rate for each dominance pairing, computed from linear combinations of the main effect of tie strength and the relevant interaction terms. Negative slopes indicate that stronger ties are associated with lower call rates. (**c**) Differences in slopes between we subtracted the call rate in the stimulus and recovery phases from the call rate in the control phaseings. Δ (mean) represents the average difference in slope between the two dyads (positive values indicate that the first dyad has the steeper slope). Associated 95% CI, posterior probabilities (Pr(β>0), Pr(β<0)), and parametric two-sided bootstrap p-values (p_boot) are shown.

**Figure 3:**
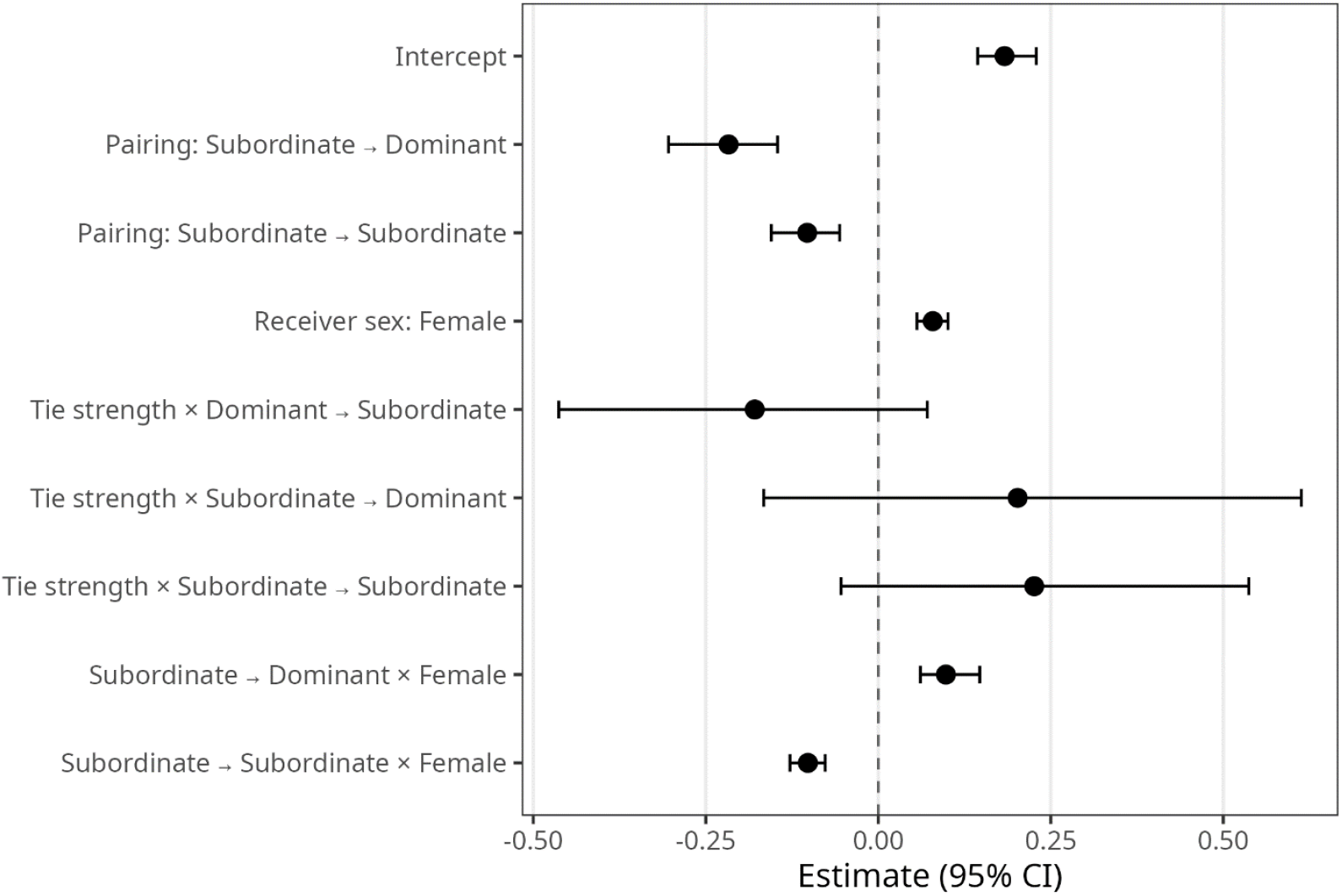
Forest plot showing estimates from the multiple imputation model testing the effect of dyadic tie strength, dominance and receiver sex on the sunning call playback reply call rate. Points represent mean parameter estimates across imputations; horizontal bars show 95% intervals. The dashed vertical line at zero indicates no effect.

**Figure 4:**
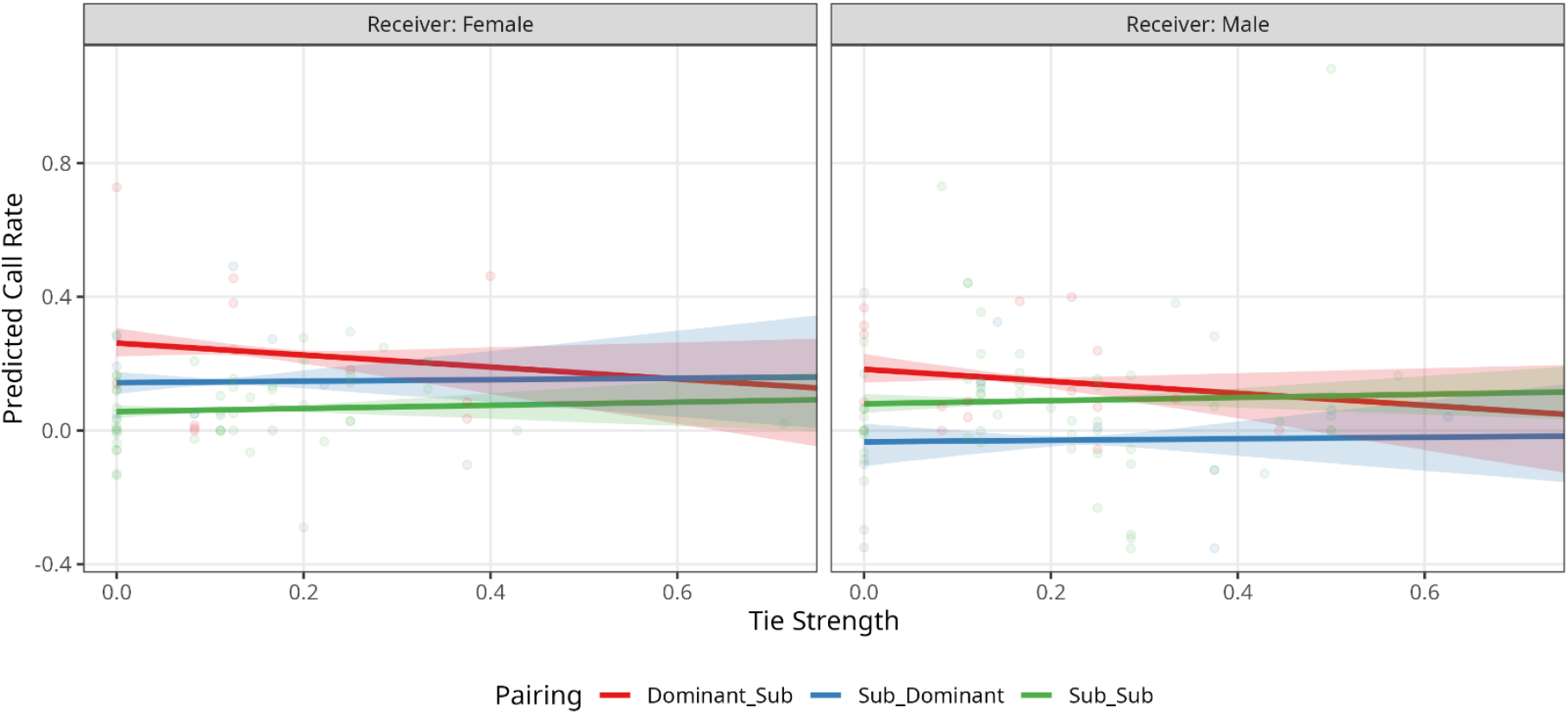
The effect of tie strength on the vocal response to sunning calls playbacks. A change in the call rate between the control and stimulus phases. The curves represent the model prediction regression with a 95% CI as shaded areas of the respective colour. The colours of the curves show the dominance relationship of sender-receiver pairs. Dominant→Subordinate (red) represents cases in which calls of a dominant sender were played to a subordinate receiver. The other two categories follow the same logic.

## DISCUSSION

The younger subordinate age classes (< 2 years) in meerkat groups showed strong within-age class association while sunning, and were also strongly associated with the dominant individuals. Adult subordinate meerkats (> 2 years), on the other hand, showed an overall weaker association within and between the age classes. Dominance, receiver sex and dyadic tie strength demonstrated a complex role in meerkat sunning call turn-taking exchanges between non-juvenile individuals (> 6 months). Overall, subordinate receivers increased their call rate when responding to dominants relative to their interactions with other subordinates.

The effect of tie strength on calling behaviour was mixed. In subordinate–dominant dyads, we observed a trend towards higher response rates when ties were weaker. By contrast, in subordinate–subordinate and subordinate-dominant dyads, tie strength had essentially no effect on calling behaviour. These across dyad differences were estimated with wide uncertainty and should be interpreted cautiously. Thus, only part of our predictions about tie-strength effects were supported, and further data will be needed to assess whether subordinate investment in weak bonds with dominants is a robust pattern. Generally, meerkats showed an increase in sunning call rate during playbacks, which independently replicates our previous results showing that meerkat sunning calls are socially facilitated as part of vocal interactions with conspecific individuals (Demartsev et al., 2018). Furthermore, females showed an overall higher vocal participation in sunning call interactions in comparison to males.

A stronger response to the sunning calls of dominants is consistent with a potential social regulatory role and aligns with earlier suggestions that sunning calls may have appeasement functions (Demartsev et al., 2018). The despotic structure of meerkat groups has strong implications for the social dynamics between subordinate and dominant individuals. Aggressive dominance assertions by dominant females, to the extent of full eviction of subordinate females from the group, are common in the later stages of the dominant female’s pregnancy (Kutsukake and Clutton-Brock, 2006a). Evicted and nearly-evicted individuals often attempt to re-integrate into the group by exhibiting submissive behaviour towards the dominants (Clutton-Brock et al., 2010; Reber et al., 2013) or undertaking “pay-to-stay” tactics (e.g. providing alloparental care, MacLeod et al., 2013).

Dominance-related aggression has a long-term negative effect on the relationship between opponents in meerkats, and only avoidance has been reported to effectively reduce received aggression, as reconciliation behaviour has not been observed, and submission did not appear to affect aggression rates (Kutsukake and Clutton-Brock, 2008a). In Dominant-Subordinate interaction dyads, the receivers’ call rate increased, possibly further modulated by the dyadic tie strength, highlighting the importance of dominance in meerkat groups. The value of having stable bonds with the dominant individuals has been suggested to drive subordinate meerkats to invest in grooming (Kutsukake and Clutton-Brock, 2006b; Kutsukake and Clutton-Brock, 2010) and submission behaviours, especially when more vulnerable (Reber et al., 2013). Sunning call exchange patterns could have similar drivers, as coordinated vocal exchanges are often attributed to a prosocial function, reducing tension and building social relationships (Fedurek et al., 2013; Laporte and Zuberbühler, 2010).

Vocal signals towards dominant individuals have been associated with submission (Kutsukake and Clutton-Brock, 2008a), reconciliation (Silk et al., 1996) and willingness to interact (Laporte and Zuberbühler, 2010). Submission behaviour in meerkats is often accompanied by high-pitched vocalisations and a crouching posture (Kutsukake and Clutton-Brock, 2006a) and while sunning calls are similar in acoustic structure to submission calls, sunning individuals do not assume a crouching posture and the calls are produced by most individuals in the group, regardless of their dominance status or sex (Demartsev et al., 2018). Thus, the overall low conflict context of sunning behaviour in meerkats makes it less likely that sunning call exchanges serve as strictly submission. Reconciliation, if defined broadly as improving weakened social bonds (Aureli, 1997), rather than an immediate and specific set of post-conflict behaviours, could also potentially fit the observed patterns of subordinates increasing their call rate towards dominant individuals. However, the higher call rates of dominant females compared to dominant males, when interacting with subordinates, are harder to explain in the context of reconciliation.

Alternatively, increased vocal activity when interacting with dominants could be related to heightened stress (Lemasson et al., 2010). In red-capped mangabeys (*Cercocebus torquatus*), large hierarchical gaps between callers have been shown to cause shorter response times in both the high-ranking and low-ranking partners (Meunier et al., 2023). Speaking tempo in humans has been shown to increase under both stressful (Kirchhübel et al., 2011) and joyful (Scherer, 2003) conditions suggesting that arousal rather than valence (Briefer, 2012) might be driving the increase in vocal activity. Similarly, stress response alone would not explain the divergent call rates between males and females across different dominance dyads. The changes in call rate are likely determined by a more complex motivation when interacting with socially valuable or influential group members.

Engaging in continuous and reciprocal interaction, where each partner’s behaviour influences the next, has been linked to cooperation (Taborsky and Riebli, 2020; Takahashi et al., 2013) and tolerance (Smith et al., 2011). For a subordinate meerkat, improving a bond with an influential dominant is crucial. While our results provide only circumstantial evidence for this pattern, investing in a pro-social interaction with dominant individuals would fit the “vocal grooming” functionality. Within the subordinate class, rare aggression and a lack of steep hierarchies (Kutsukake and Clutton-Brock, 2006a), may lead to weak drivers towards improving bonds with other subordinates and a general preference for interacting with already bonded partners. It is essential to acknowledge that our chosen social context (proximity while sunning) offers only a snapshot of the complex social relationships of meerkat groups. Future work incorporating multilayer network approaches that combine multiple affiliative behaviours would provide a more nuanced test of the role of vocal exchanges in relationship management (Finn et al., 2019).

It is evident that the observed response patterns further diverge across the sexes. During the breeding season, as the reproductive conflict between the dominant and subordinate meerkats intensifies (Young et al., 2006) these contrasts could become starker. Our sample size did not allow us to robustly test the differences between all possible combinations of sex, dominance and bond strengths of both sender and receiver individuals. However, while reproductive conflict clearly affects the social relationships of group members (Kutsukake and Clutton-Brock, 2008b), meerkat groups ultimately comprise a cohesive mixed-sex social unit with individuals of both sexes participating in cooperative activities i.e. sentinel (Rauber and Manser, 2018), babysitting (Clutton-Brock et al., 2000), pup feeding (Carlson et al., 2006), and mobbing (Graw and Manser, 2007). Thus, bond maintenance and conflict prevention are not limited to within-sex mechanisms and are likely to also act more generally to support the social structure and functionality of a group. And manifest outside of the breeding season.

The tendency for individuals to interact more with closely (Kulahci et al., 2015) or loosely bonded partners (Chereskin et al., 2022) can have different underlying causes. Maintenance of stable bonds requires effort, and vocal grooming may function as an addition to physical grooming (Kulahci et al., 2015). In comparison, the relatively low cost of vocal exchanges could be beneficial for extending individuals` social connectivity and maintaining valuable but loose affiliations (Chereskin et al., 2022). Similarly, vocal exchanges could be used for establishing novel ties, in a process similar to vocally mediated pair formation in songbirds (D’Amelio et al., 2017). Priming (establishing) ties remotely would be especially useful when physical proximity between weakly bonded individuals might result in aggressive interactions, as could be the case between dominant and subordinate meerkats in some cases.

While this work has demonstrated a correlation between vocal exchanges and social relationships, we have yet to establish a causal link between these variables. In addition, the mechanism by which such a causal relationship might act remains unclear. Given that cooperative activities are often associated with the strengthening of bonds in animals, including humans (Cheney, 2011; Heesen et al., 2020), one possibility is that coordinated signalling exchanges could serve as a ritualised version of joint action, signifying commitment to the partner and the task (Heesen et al., 2020; Wolf et al., 2016). Across taxa, such interactions can contribute to the establishment of new relationships, for example, in the pair formation context of duetting in birds (Hall, 2004) or gibbons (Geissmann and Orgeldinger, 2000) as well as to the reinforcement of existing ones. Continuous and repeated call exchanges could be a form of cooperation (Pougnault et al., 2022), stabilising relationship, increasing social tolerance (Cheney et al., 1995) and cohesion.

The accumulation of potential examples of vocal grooming beyond the primate lineage suggests that it could be a common phenomenon in nature. Our findings provide evidence that sunning calls interaction reflects dominance and possibly finer social relationships in meerkat groups. The skew towards stronger responses to dominants provides a preliminary indication of the social regulation function of these calls and contributes to comparative work on “vocal grooming” across taxa. While our study does not establish a causal link, it highlights testable pathways by which vocal exchanges could function in bond reinforcement or the management of weak but socially valuable ties. More broadly, the idea that low-cost communicative interactions can support cohesion has been proposed as relevant to the evolution of human sociality and language (Dunbar, 1998; Kulahci et al., 2015). Future longitudinal studies that track both relationship quality and vocal dynamics will be essential for evaluating these functional hypotheses and for clarifying the generality of vocal grooming across species.

