## Supplementary material for "Dominance asymmetries shape vocal exchanges in meerkats": Supp

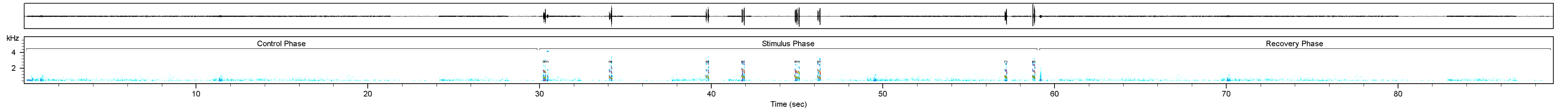
kHz
4
2
Control Phase
Stimulus Phase
Recovery Phase
10
20
30
40
50
60
70
80
Time (sec)

| Year | Group # | Total individuals | Non-Juv | Juveniles | Group Code |
|---|---|---|---|---|---|
| 2019 | 1 | 13 | 11 | 2 | W |
| 2019 | 2 | 19 | 13 | 6 | JX |
| 2019 | 3 | 15 | 10 | 5 | HM |
| 2019 | 4 | 18 | 13 | 5 | L |
| 2021 | 5 | 15 | 10 | 5 | L |
| 2021 | 6 | 22 | 16 | 6 | MP |
| 2021 | 7 | 13 | 13 | 0 | ZU |

**Table S1**: Composition of of the study groups. Juveniles are group members that were younger than 6 month. Non-Juvenile individuals are all group members that were older than 6 months at the time of the study
